# RExPRT: a machine learning tool to predict pathogenicity of tandem repeat loci

**DOI:** 10.1101/2023.03.22.533484

**Authors:** Sarah Fazal, Matt C. Danzi, Isaac Xu, Shilpa Nadimpalli Kobren, Shamil Sunyaev, Chloe Reuter, Shruti Marwaha, Matthew Wheeler, Egor Dolzhenko, Francesca Lucas, Stefan Wuchty, Mustafa Tekin, Stephan Züchner, Vanessa Aguiar-Pulido

## Abstract

Tandem repeats (TRs) are polymorphic sequences of DNA that are composed of repeating units of motifs, whose lengths can vary depending on the type of TR. Expansions of TRs are responsible for approximately 50 monogenic diseases, compared to over 4,300 disease causing genes disrupted by single nucleotide variants and small indels. It appears thus reasonable to expect the discovery of additional pathogenic repeat expansions, which has the potential of significantly narrowing the current diagnostic gap in many diseases. Recently, short and long-read whole genome sequencing with the use of advanced bioinformatics tools, have identified a growing number of TR expansions in the human population. The majority of these loci are expanded in <1% of genomes. Categorizing and prioritizing such TR loci is a growing challenge to human genomic studies. We present a first-in-class machine learning tool, RExPRT (Repeat EXpansion Pathogenicity pRediction Tool), which is designed to distinguish pathogenic from benign TR expansions. Leave-one-out cross validation results demonstrated that an ensemble approach comprised of support vector machines (SVM) and extreme gradient boosted decision tree (XGB) classify TRs with a precision of 92% and a recall of 90%. Further validation of RExPRT on unseen test data demonstrate a similar precision of 86%, and a recall of 60%. RExPRT’s high precision in particular, will be of significant value to large-scale discovery studies, which require the prioritization of promising candidate loci for time-consuming and costly functional follow-up studies. Application of RExPRT to ~800,000 TRs in the reference genome identified ~30,000 TRs that would be likely pathogenic upon expansion. Thus, RExPRT establishes a foundation for the application of machine learning approaches to categorize the pathogenicity of tandem repeat loci.

## Introduction

Tandem repeats (TRs) are regions of the DNA that are composed of repeating motifs that vary between 2 - 6 base pairs (bp) in length.^1^ There are 1.5 million TR loci scattered throughout the human genome.^2^ Expansions of TRs can produce changes in the underlying genetic architecture, thus impacting molecular processes through the RNA or protein level.^3^ Currently only ~50 known disease-causing tandem repeat expansion loci have been identified,^3^ a small minority compared to the 4,361 genes associated with Mendelian diseases (OMIM).^4^ We speculate that there are many more disease-associated TR loci to be discovered, and progress has been hindered by the technical challenges associated with identifying long expanded repeats from short-read sequencing data. Specifically, correctly aligning short reads containing fully repetitive sequence, and determining repeat lengths based on incomplete sequence information has posed significant complications. However, with the advancement of tools such as ExpansionHunter^5,6^ and GangSTR^7^, as well as the emergence of databases characterizing tandem repeats (TRs) in control samples,^8^ the identification of rare repeat expansions from patient genomes is now possible. Previously, we showed that genomes from healthy individuals each have an average of ~250 common large TRs (> 175bp) and a median of three rare large TRs.^8^ This indicates that rare repeat expansions can be benign and population allele frequency alone cannot reliably distinguish pathogenic loci.

To assess the pathogenicity of single nucleotide variants (SNVs) and small indels, detailed guidelines assist with filtering variants that go beyond consideration of allele frequencies.^9,10^ Tools such as ANNOVAR^11^ and Ensembl’s Variant Effect Predictor (VEP)^12^ annotate variants with their functional impact on the corresponding protein to assist with such filtering. The commonly-used pathogencity predictor Combined Annotation-Dependent Depletion (CADD) scores simple variants based on sequence context including evolutionary constraint, epigenetics, and gene model annotations.^13^ Numerous additional tools are available for variant annotation, prediction, and prioritization in SNVs and small indels.

For structural variants (SVs), several tools have recently been developed to aid in variant prioritization. Specifically focusing on structural variants in exons, StrVCTVRE uses supervised learning to distinguish pathogenic from benign SVs, accounting for features such as conservation, expression, and exon structure.^14^ Another tool, SVpath, predicts the pathogenicity of exonic SVs by incorporating features that are based on functional impact scores of overlapping SNVs, as well as gene level and transcriptomics scores.^15^ Developing this approach further, DeepSVP integrates ranking of noncoding variants and incorporates phenotype information using a deep learning approach to improve the selection of patient-specific variants in a more precise manner.^16^

In the field of TRs, efforts to prioritize variants have focused on an underlying assumption that TR constraint correlates with pathogenicity. Gymrek *et al*. showed this to be true for select early-onset disease loci such as RUNX2 and HOXD13.^17^ However, according to their methodology for predicting mutational constraint, late onset disease loci such as ATXN7 are not highly constrained. Since many repeat expansion diseases manifest later in life, mutational constraint alone is an unreliable prioritization metric. One of the few existing tools for TR prioritization – SISTR - is based on a population genetics framework.^18^ It calculates a selection coefficient, which incorporates measures of mutation, genetic drift, and negative natural selection. While alleles that are negatively selected are predicted to be more deleterious, these values correlate with mutational constraint and are therefore subject to the same limitations. Additionally, SISTR was demonstrated for use in a complex genetic disorder, autism, whose underlying genetic etiolgy differs from those of rare Mendelian diseases.^18^

Currently, there are no prioritzation models built on labelled training data containing examples of both pathogenic and benign TRs. The challenge lies especially in the limited training data, considering there are only ~50 known pathogenic repeat expansion loci discovered to date.^3^ To address this gap, we present RExPRT, a supervised machine learning-based **R**epeat **Ex**pansion **P**athogenicity p**R**ediction **T**ool. RExPRT is the first tool applicable for both early and late onset TR-driven rare monogenic diseases that can score and categorize TR loci as pathogenic and benign.

## Materials and Methods

### Identification of significant data features

To select features that may be relevant for classifying TR loci as pathogenic or benign, we downloaded 30 different annotation datasets from a variety of sources including UCSC’s Table Browser, PsychENCODE, and relevant publications (Supplementary Table 1). We converted all dataset files into BED format. First, we created a list of 40 known pathogenic TRs which are listed as represented in the training dataset in Table 1. These TRs were included in initial analyses because they are well established in their causal link to disease.^19–21^ Next, we obtained a dataset of all TRs cataloged in the reference genome from UCSC’s Table Browser (Simple Repeats table in hg19). Finally, we used this reference TR dataset to create a set of “matched TRs”, which included all TRs that have a disease motif and occur in the same genic region. For example, all TRs with a CAG motif that occur in exons would be part of the matched TRs dataset. The three TR datasets (pathogenic TRs, reference TRs, and matched TRs) were then intersected individually with each of the 30 annotation files. Using bedtools fisher we ran Fisher’s exact tests to provide an odds ratio, indicating if the amount of overlap between pairs of TR and annotation files is statistically significant given their coverage and the size of the genome. Using these contingency tables we calculated 95% confidence intervals for the odds ratios using fisher.test() from the R stats package. If the 95% confidence interval of the odds ratio for an annotation’s intersection with pathogenic TRs was higher than the 95% confidence interval for the matched TRs, the annotation feature was considered significantly enriched for pathogenic TRs. For genic annotation features (3’UTR, 5’UTR, exon, intron, and promoter), the comparison was made between pathogenic TRs and reference TRs instead to avoid bias, since matched TRs already selected for certain genic regions.

**Table 1.** Pathogenic TRs used in training/LOOCV and testing datasets. RExPRT ensemble classification, as well as both SVM and XGB scores are provided for each pathogenic TR. Additionally information is provided for each locus including motif, the gene region in which the TR is located, inheritance pattern of the associated disease, whether a motif change is required to pathogenic, and whether the motif was uniquely represented among other pathogenic TRs in the training dataset.

| Gene | Motif | Gene region | Inheritance | Motif change | Unique motif | SVM score | XGB score | RExPRT classification | Dataset |
| --- | --- | --- | --- | --- | --- | --- | --- | --- | --- |
| AFF2 | CCG | 5'UTR | X-Linked | No | No | 0.99 | 0.91 | Pathogenic | Training |
| AFF3 | CGG | Intron | Dominant | No | No | 0.98 | 0.64 | Pathogenic | Testing |
| AR | CAG | Exon | X-Linked | No | No | 0.97 | 1.00 | Pathogenic | Training |
| ARX | GCG | Exon | X-Linked | No | No | 0.99 | 0.97 | Pathogenic | Training |
| ATN1 | CAG | Exon | Dominant | No | No | 0.97 | 0.99 | Pathogenic | Training |
| ATXN1 | CAG | Exon | Dominant | No | No | 0.97 | 0.99 | Pathogenic | Training |
| ATXN10 | ATTCT | Intron | Dominant | No | Yes | 0.00 | 0.00 | Benign | Training |
| ATXN2 | CAG | Exon | Dominant | No | No | 0.87 | 1.00 | Pathogenic | Training |
| ATXN3 | CAG | Exon | Dominant | No | No | 0.63 | 1.00 | Pathogenic | Training |
| ATXN7 | CAG | Exon | Dominant | No | No | 1.00 | 1.00 | Pathogenic | Training |
| ATXN8OS | CTG | 3'UTR | Dominant | No | No | 0.99 | 0.90 | Pathogenic | Training |
| BEAN1 | TGGAA | Intron | Dominant | Yes | Yes | 0.51 | 0.04 | Pathogenic | Training |
| C9orf72 | GGGGCC | Intron | Dominant | No | Yes | 0.90 | 0.94 | Pathogenic | Training |
| CACNA1A | CAG | Exon | Dominant | No | No | 0.92 | 1.00 | Pathogenic | Training |
| CBL | CGG | 5'UTR | Dominant | No | No | 0.98 | 0.98 | Pathogenic | Testing |
| CNBP | CCTG | Intron | Dominant | No | Yes | 0.07 | 0.00 | Benign | Training |
| CSTB | CCCCGCCCCGCG | 5'UTR | Recessive | No | Yes | 0.83 | 0.52 | Pathogenic | Training |
| DAB1 | TTTCA | Intron | Dominant | Yes | No | 0.00 | 0.53 | Pathogenic | Training |
| DIP2B | CGG | 5'UTR | X-Linked | No | No | 0.97 | 1.00 | Pathogenic | Testing |
| DMPK | CTG | 3'UTR | Dominant | No | No | 0.76 | 0.98 | Pathogenic | Training |
| FMR1 | CGG | 5'UTR | X-Linked | No | No | 0.97 | 0.99 | Pathogenic | Training |
| FOXL2 | GCC | Exon | Dominant | No | No | 0.99 | 0.99 | Pathogenic | Training |
| FXN | GAA | Intron | Recessive | No | Yes | 0.00 | 0.01 | Benign | Training |
| GIPC1 | CGG | 5'UTR | Dominant | No | No | 0.95 | 0.99 | Pathogenic | Training |
| GLS | GCA | 5'UTR | Recessive | No | No | 0.93 | 1.00 | Pathogenic | Training |
| HOXA13 | GCG | Exon | Dominant | No | No | 1.00 | 0.99 | Pathogenic | Training |
| HOXD13 | GCA | Exon | Dominant | No | No | 0.83 | 1.00 | Pathogenic | Training |
| HTT | CAG | Exon | Dominant | No | No | 0.97 | 1.00 | Pathogenic | Training |
| JPH3 | CTG | 3'UTR | Dominant | No | No | 0.56 | 0.62 | Pathogenic | Training |
| LOC642361 | CGG | Exon | Dominant | No | No | 0.83 | 0.04 | Pathogenic | Testing |
| LRP12 | CGG | 5'UTR | Dominant | No | No | 0.98 | 0.99 | Pathogenic | Training |
| MARCHF6 | TTTCA | Intron | Dominant | Yes | No | 0.00 | 0.01 | Benign | Testing |
| NBPF8 | GGC | 5'UTR | Dominant | No | No | 0.88 | 0.71 | Pathogenic | Training |
| NOP56 | GGCCTG | Intron | Dominant | No | Yes | 0.67 | 0.66 | Pathogenic | Training |
| PABPN1 | GCG | Exon | Dominant | No | No | 0.90 | 0.80 | Pathogenic | Training |
| PHOX2B | GCA | Exon | Dominant | No | No | 0.71 | 0.99 | Pathogenic | Training |
| PPP2R2B | CAG | 5'UTR | Dominant | No | No | 0.87 | 1.00 | Pathogenic | Training |
| RAPGEF2 | TTTCA | Intron | Dominant | Yes | No | 0.02 | 0.03 | Benign | Testing |
| RFC1 | AAGGG | Intron | Recessive | Yes | Yes | 0.01 | 0.00 | Benign | Training |
| RUNX2 | GCG | Exon | Dominant | No | No | 1.00 | 0.99 | Pathogenic | Training |
| SAMD12 | TTTCA | Intron | Dominant | Yes | No | 0.94 | 0.00 | Pathogenic | Training |
| SOX3 | GCC | Exon | X-Linked | No | No | 0.95 | 0.98 | Pathogenic | Training |
| STARD7 | TTTCA | Intron | Dominant | Yes | No | 0.05 | 0.49 | Benign | Testing |
| TBP | CAG | Exon | Dominant | No | No | 0.98 | 1.00 | Pathogenic | Training |
| TCF4 | CTG | Intron | Dominant | No | No | 0.62 | 0.96 | Pathogenic | Training |
| TNRC6A | TTTCA | Intron | Dominant | Yes | No | 0.00 | 0.02 | Benign | Testing |
| **XYLT1** | GGC | 5'UTR | Recessive | No | No | 0.99 | 0.04 | Pathogenic | Training |
| **YEATS2** | TTTCA | Intron | Dominant | Yes | No | 0.02 | 0.63 | Pathogenic | Testing |
| **ZIC2** | GCT | Exon | Dominant | No | No | 0.98 | 1.00 | Pathogenic | Training |
| **ZNF713** | CGG | 5'UTR | Dominant | No | No | 0.97 | 0.99 | Pathogenic | Testing |

### Quantitative and multivalued features

We further selected additional annotations to test in our models. These included datasets with numerical values rather than the purely categorical datasets discussed above. For the features described above, TRs were given a value of 1 if the locus intersected with the annotation feature, and a value of 0 if there was no intersection. Quantitative datasets that we incorporated include pLI scores, LOEUF scores,^22^ and GERP scores.^23^ The basis for including these is due to their significance in assessing pathogenicity of SNVs. We also created a feature that calculated the distance between the TR locus and the nearest gene, which was important for TRs that are intergenic.

Additionally, we incorporated information regarding the motif assuming a motif-phenotype correlation exists as proposed by Ishiura et al, where TRs with the same motifs, occurring in the same genic regions - albeit in different genes entirely - can produce the same phenotypes, suggesting TR motif may be important in its pathogenicity.^24^ We calculated the percentages of each nucleotide in the motif and provided a GC content as well. For motif analysis we also used the Sequence to Star Network (S2SNet) approach,^25^ which can transform any character-based sequence into a graph-based star-shaped complex network. We characterized the star network’s topological indices (Tis) with calculations of different metrics, including its Shannon entropy, spectral moments, Harray number, Wiener index, Gutman topological index, Schultz topological index, Balaban distance connectivity index, Kier-Hall connectivity index, and Randic connectivity index.^25^ S2SNet was downloaded from GitHub and run using the default parameters on the command line. For the input, we created a sequence of 10 repeating units of the TR motif.

Finally, we accounted for tissue expression of the genes that harbored TRs. We created a categorical feature reflecting the tissue where each gene had its maximum expression according to data from the GTEx Portal (date of accession: September 2021). The TR was then assigned to one of three possible categories: expressed in neurological tissue, expressed in another, or unknown expression.

### Training and testing dataset creation and preprocessing

To train RExPRT, we used the same 40 known pathogenic loci that were assessed with Fisher’s exact tests above. Based on our previous work,^8^ we used a set of 754 TR loci that were commonly expanded in the 1000 Genomes Project controls as negative training data. Since these TRs are commonly expanded, we presumed that they are benign. To annotate our TRs with the features discussed above, we created an input file containing the reference coordinates of the locus in hg19 (chromosome, start position, and end position), as well as the motif. After annotation with the features described, we used the_OneHotEncoder_ from Scikit-Learn to preprocess the data. This was used for categorical variables including GTEx, gene region (intron/exon/intergenic), as well as gene location (first/middle/last).

For the testing dataset, we used the 10 remaining pathogenic TR loci listed in Table 1. These were loci that were either discovered more recently or have only been found in a small number of affected families.^3^ For the negative loci in the test set, we used sites that can be presumed benign, but are still rare since RExPRT will have to distinguish between rare benign TRs and rare pathogenic TRs. We decided to use 83 rare TRs that were candidate TRs resulting from running the outlier pipeline described subsequently on our 102 positive controls. Since these genomes correspond to patients that were all diagnosed with a Mendelian repeat expansion disorder (caused by a single gene), other TR expansions in these genomes are presumed to be benign. Therefore, we combined all the candidate TRs from these genomes and excluded all known pathogenic TRs to produce a list of 83 rare, benign TRs.

### Model testing and assessment

We tested seven different statistical learning models: logistic regression, k-nearest neighbors (KNN), support vector machine (SVM), random forest (RF), decision tree (DT), gradient boosted decision tree (XGB), and linear discriminant analysis (LDA). To evaluate the performance of these models, we used a leave-one-out cross validation (LOOCV) approach. The LOOCV approach is useful here because we have a small positive training dataset of known pathogenic TRs. For models that are sensitive to scaling such as SVM, we used _StandardScaler_ to standardize features by subtracting the mean and scaling by the standard deviation. To assess model performance, we obtained the confusion matrix and determined overall accuracy, precision, recall, and F1 metrics. We also produced receiver operating characteristic curve (ROC) and precision-recall curves (PRC) and calculated the area under the curve for each one. All coding was done in Python using Scikit-Learn. Formulas for calculations are listed below:

1. 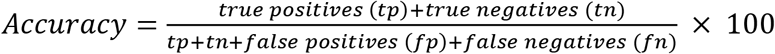
2. 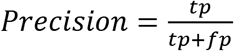
3. 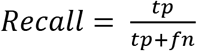
4. 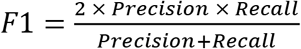

### Feature selection

To further improve the accuracy of the two best performing models, we plotted a feature importance graph (XGB) and a permutation importance graph (SVM) to visualize the significance of each feature to the models. We then removed features that were ranked low on the list, indicating that the feature was not adding significant value in the decisionmaking process. We only removed features if they did not impact confusion matrix scores or the overall area under the ROC curve (AUROC) score.

Additionally, we created a correlation plot of all features and removed features that were highly correlated. In particular, correlated features that produced the highest AUROC value if retained were kept, ensuring that removal of features did not reduce the AUROC score.

### Hyperparameter tuning

We used Scikit-Learn’s GridSearchCV to fine-tune the SVM and XGB models. For SVM we tuned three parameters: C, gamma, and kernel. For XGB we tuned two parameters: number of estimators, and max depth.

### Ensemble method

To improve the overall recall of RExPRT, we devised an ensemble approach, which encompasses the two different models described above (SVM and XGB).^26^ The classification of the TR by the ensemble model will be pathogenic if either of the models predict likelihood of pathogenicity. The threshold is set to a standard 0.5 probability, above which the TR is classified as pathogenic. Furthermore, RExPRT also provides a confidence score, which is the sum of the SVM and XGB scores. Figure 1 is a schematic illustrating the complete methodology and workflow of RExPRT, covering training, LOOCV, model selection, and predictions on the testing dataset.

**Figure 1.**
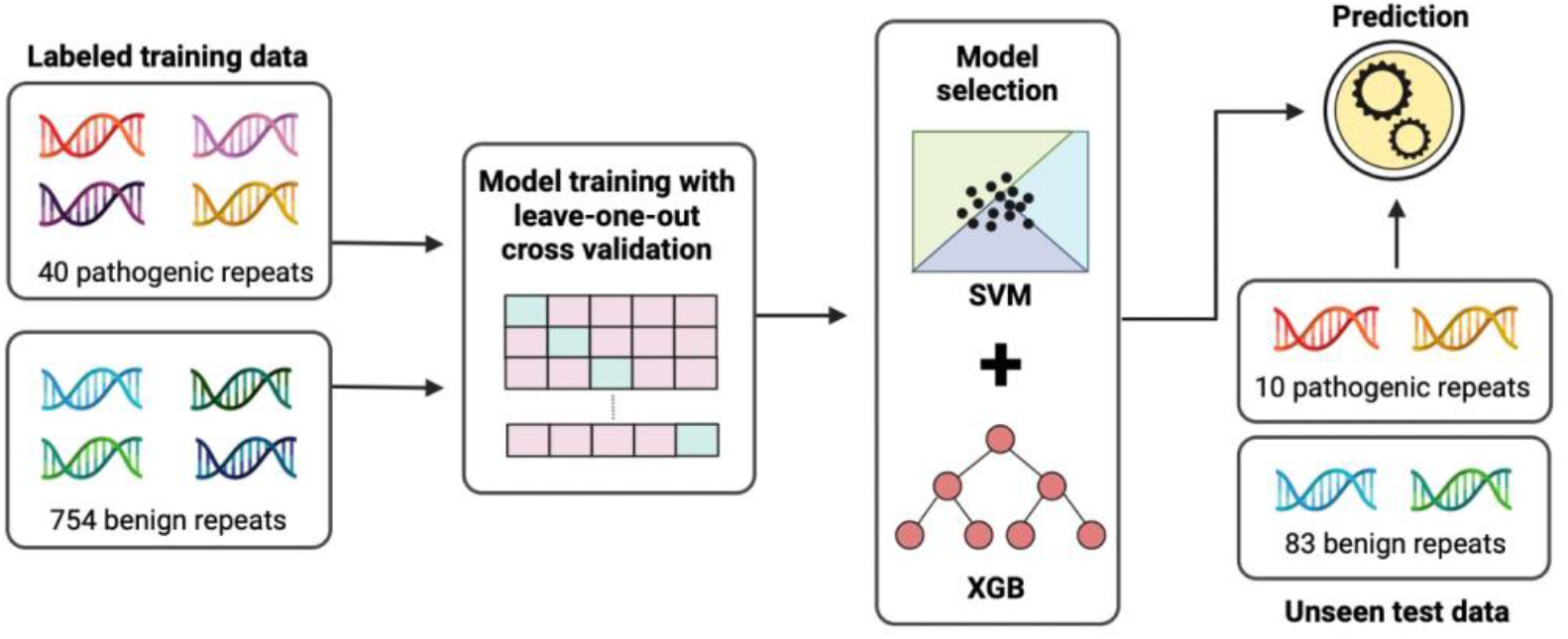
Methodology of RExPRT. RExPRT was trained on 40 known pathogenic TRs and 745 benign TRs that are commonly expanded in the 1000 Genomes Project controls. These TRs were annotated with 46 features, which are used in a supervised statistical learning approach to classify TRs as pathogenic or benign. Seven different models were trained and validated using the LOOCV technique. Two models were selected and fine-tuned to create an optimized ensemble method for ranking repeats. Ten pathogenic TRs and 83 rare, benign TRs were used for testing RExPRT’s performance.

### *Analysis of* ~800,000 *reference repeats classified by RExPRT*

We applied RExPRT to *836,545* TRs with motif lengths between 3 and 8bp listed in the hg19 reference genome. Bedtools fisher was used to perform Fisher’s exact tests and calculate odds ratios for gene regions and regulatory regions. For gene expression characterization, we excluded intergenic TRs. We then performed Fisher’s exact tests in R, first creating a contingency table for benign and pathogenic TRs. To do this, we calculated the number of genes that overlapped in the different sets of tissues with the two groups of classified TRs.

For analysis of Online Mendelian Inheritance in Man (OMIM) genes, we downloaded the list of genes from their website, and filtered for those that are Mendelian. For those genes that are dominant and recessive, we used this same dataset and filtered for the respective subtype. The ataxia gene list was obtained from GeneDx’s Ataxia Xpanded panel (https://www.genedx.com/tests/detail/ataxia-xpanded-panel-887). The odds ratios were obtained by calculating Fisher’s exact tests in R, as described for the gene expression data.

To analyze the disease motifs, we filtered all pathogenic and benign TRs for those which contained a known repeat expansion disease motif (Table 1). We included each window shift of the motif as well as its reverse complement. For pure GC motifs, we filtered for TRs that only contained G and C in their motif. For polyglutamine (polyQ) and polyalanine (polyA) motifs, we began with only coding TRs in each group and filtered for the trinucleotides which code for these amino acids. Since we do not have the coding frame information, many of these will not actually code for glutamine or alanine, so this is an overestimation. Odds ratios were calculated in R, as described for the gene expression, and disease genes categories.

### Undiagnosed Diseases Network (UDN) sample processing

Patient fastq files were aligned with burrows wheeler aligner (BWA) to the GRCh38 reference genome.^27^ The resulting SAM files were converted to BAM files. Duplicates were removed with Picard tools MarkDuplicates - https://github.com/broadinstitute/picard. After sorting and reindexing, base quality score recalibration (BQSR) was performed using genome analysis toolkit (GATK).^28^ Next, ExpansionHunter Denovo (EHDn) was run on the resulting BAM files, using additional parameters ‘--min-unit-len 3 --max-unit-len 8’.^6^ The outputs of EHDn for each case was then aggregated separately to the outputs for the 2,405 samples from the 1000 Genomes Project controls using EHDn’s helper scripts to allow depth-normalization. Finally, the bed file output was expanded from its sparse encoding into a dense matrix format using R. This dense matrix of depth-normalized anchored in-repeat read (IRR) counts was used as the input for the outlier pipeline.

### Outlier pipeline

After sample processing, each case is represented by a dense matrix of TRs with anchored IRR counts for the case as well as all controls. Summary statistics were calculated for all TRs before filtering. For each TR, the outlier pipeline calculates a z-score, a kernel density estimation, and the percentage of controls with anchored IRR counts above that seen for the patient sample. The z-score for a case is a measure of their anchored IRR count in terms of the number of standard deviations above or below the mean anchored IRR counts observed in controls. The kernel density estimation gives the expected proportion of controls with anchored IRR counts above that of the patient, based on the density curve distribution of controls. All three measures give us an indication of the likelihood of the anchored IRR count for the patient sample being part of the distribution of controls. Downstream filtering selects for TRs with high z-scores, and low values for the other two measures. Specifically, the pipeline filters out TRs based on anchored IRR counts in patients (>=5), frequency of controls with at least 1 allele over 175 bp (<1%), genomic region the TR is located in (≠intergenic), and whether the TR stems from a reference repeat locus or *Alu* element. Moreover, we also filter out a list of 126 false positive sites that were selected based on their presence in a heterogenous group of ~600 disease genomes and low occurance in the 1000 Genomes controls. It is thought that since these variants are too common in a cohort of rare disease cases, they cannot be causal variants. The final output is a list of candidate TRs for each patient genome. Therefore, here we define candidate TRs as TRs that are rare and expanded in a patient, occur within or close to genes, and stem from a reference repeat locus.

### Analysis of UDN genomes

We ran 2,982 genomes (968 probands and affected/unaffected family members) from the UDN through our outlier pipeline, resulting in a list of candidate TRs that can be defined as rare, genic TR expansions that stem from a reference repeat locus. Since these TRs were called by ExpansionHunter Denovo (EHDn), they do not have precise locus specificity. EHDn provides coordinates of ~1-2,000 bp surrounding the repeat. To run ExpansionHunter on these loci for determining repeat number and zygosity, we used ehdn-to-eh (https://github.com/francesca-lucas/ehdn-to-eh), which provides precise coordinates for the TR. These coordinates were then used to create the variant catalogs for running ExpansionHunter. Since the UDN genomes are aligned to hg38, we converted the loci into hg19 using UCSC’s LiftOver tool,^29^ and then ran RExPRT on the candidate TRs.

## Results

### Pathogenic TRs are enriched in regulatory regions and in specific areas within genes

We investigated whether known pathogenic TRs are enriched in any particular regions of the genome compared with matched TRs and all reference TRs. We found that pathogenic TRs are enriched in topologically associated domain (TAD) boundaries (odds ratio (OR)= 7.55; 95% confidence interval (CI) = [2.82, 17.37]), open regulatory regions (ORegAnno) (OR = 10.21; 95% CI = [5.09, 21.56]), as well as SMC3 (OR = 32.34; 95% CI = [13.54, 69.56]) and RAD21 transcription factor binding sites (OR = 37.51; 95% CI = [15.71,80.70]) (Fig. 2a). Furthermore, pathogenic TRs are also more likely to be classified as expression TRs (eTRs) (OR = 294.52, 95% CI = [99.81, 732.38]), which are repeats whose length has been reported to be directly associated with transcript levels of the respective gene.^2^

**Figure 2.**
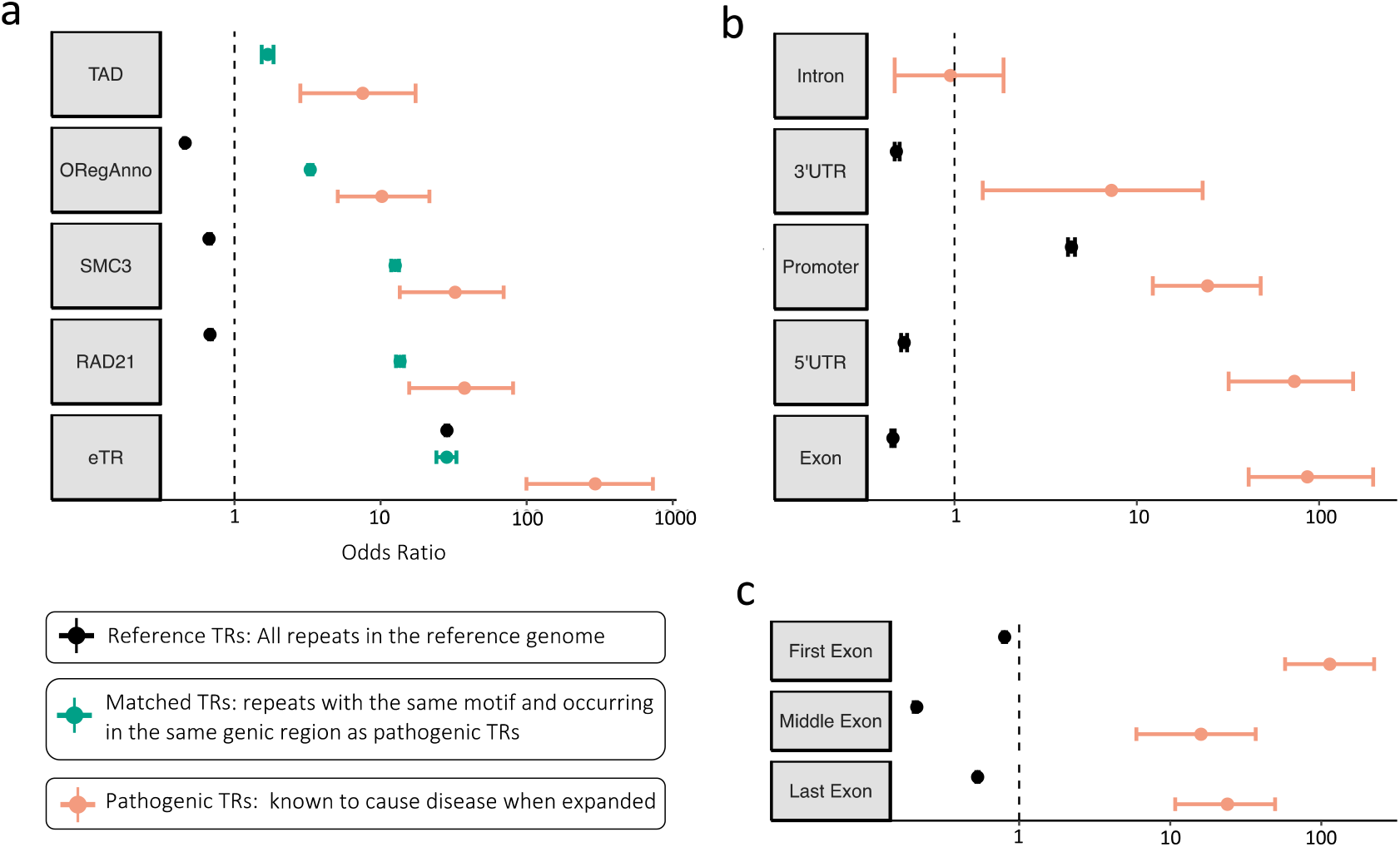
Genomic regions with enrichment of pathogenic TRs. (**a**) Odds ratios from fisher’s exact tests for pathogenic TRs, matched TRs, and reference TRs in their intersection with TAD boundaries, open regulatory regions, SMC3 and RAD21 transcription factor binding sites, and eTRs. (**b**) Odds ratios from fisher’s exact tests for pathogenic TRs and reference TRs in their intersection with different genic regions. (**c**) Odd’s ratios from fisher’s exact tests for pathogenic TRs and reference TRs in their intersection with different exonic regions.

Pathogenic TRs are also enriched in 3’UTRs (OR = 7.27; 95% CI [1.43, 22.97]), promoters (OR = 24.41; 95% CI = [12.24, 47.76]), 5’UTRs (OR = 73.20; 95% CI = [31.90, 153.71]), and exons (OR = 86.15; 95% CI = [41.10, 197.73]) while we did not find any significant enrichment in introns (OR = 0.95; 95% CI = [0.47, 1.86]) (Fig. 2b). Upon further exploration, we found that there is significantly more enrichment of pathogenic TRs in the first exon of genes (OR = 113.60; 95% CI = [57.47, 223.43]) compared to the middle and last exons (Fig. 2c).

### RExPRT achieves 99% accuracy using LOOCV

RExPRT is an ensemble method combining the two best performing models based on the LOOCV results: SVM and XGB. It excels in its low false positive rate (0.38%) while still attaining an excellent recall of 90% (Fig. 3a, b). This means we have a good balance between precision (92.31%) and recall, resulting in an F1 score of 0.91 (Fig. 3b). The corresponding ROC curve in Figure 3c highlights RExPRT’s prowess as evidenced by a high AUROC value of 0.97. While ROC curves are a standard way to present machine learning results, a PRC is more informative with an imbalanced dataset. The PRC curve is also robust (AUPRC = 0.92) and illustrates the fine balance between our precision and recall rates (Fig. 3d). Since pathogenic TRs compose [5%] of the dataset, an AUPRC value of [0.05] represents random guessing performance. Therefore, RExPRT’s performance is considerably better than random guessing. Figures 3e and f represent the results of feature or permutation importance analyses for each of the two models. We find that both models are extremely reliant on whether the TR is in an exon or 5’UTR. The SVM model utilizes pLi and LOEUF scores, and percentage calculations of nucleotides within the TR motifs. On the other hand, XGB applies the S2SNet topological indices calculations for its predictions (Fig. 3e, f). The two models complement one another, as together they boost the overall performance of RExPRT.

**Figure 3.**
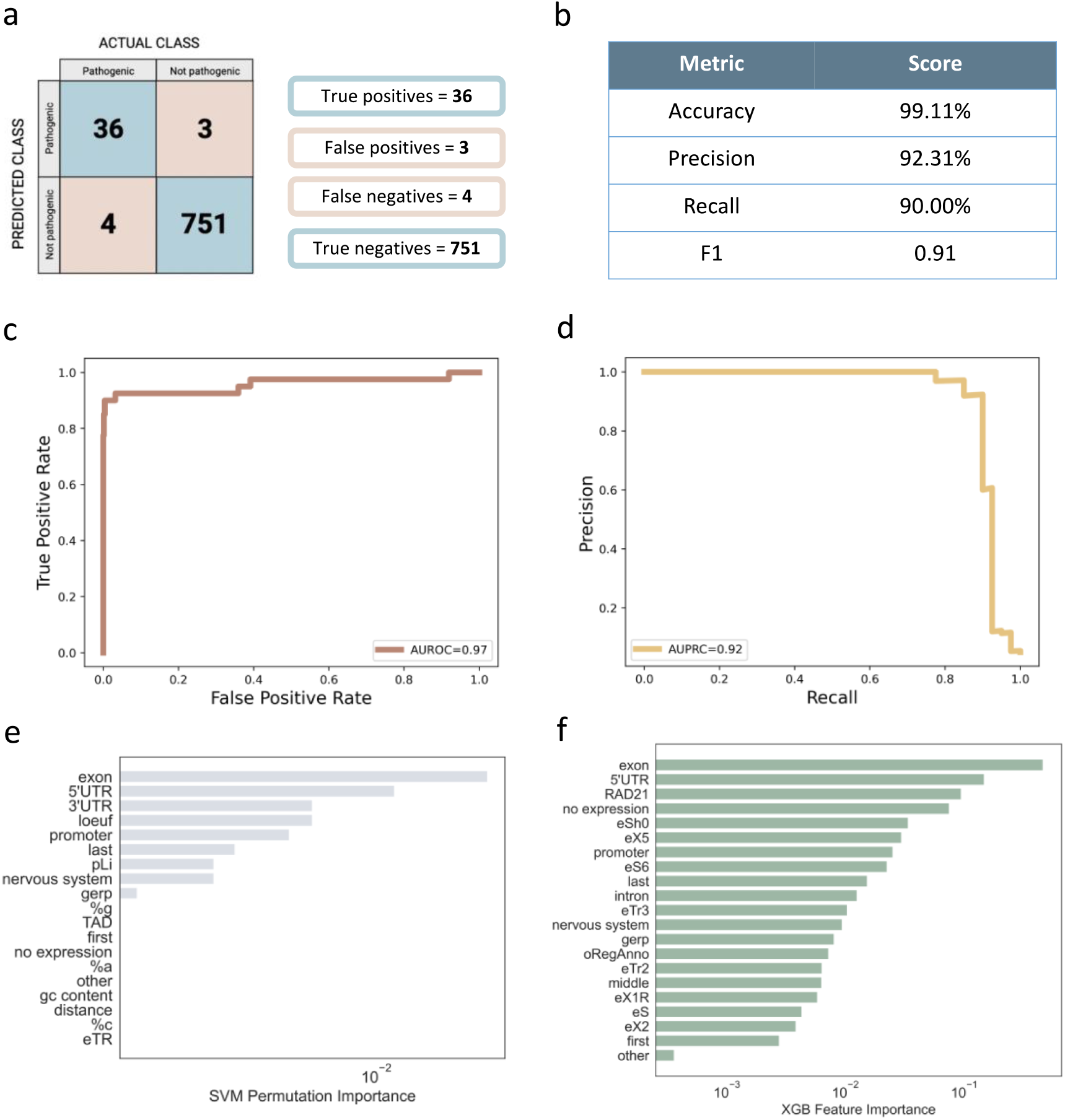
Results from ensemble of SVM and XGB with LOOCV on training dataset. (**a**) Confusion matrix outlining the number of true positives, false positives, false negatives, and true negatives on the training set. (**b**) Calculations of the accuracy, precision, recall, and F1 scores. (**c**) The ROC curve for the ensemble model and its AUC value. (**d**) PRC curve for the ensemble model and its AUC value. (**e**) Permutation importance, indicating features that allow decision making in the SVM model and (**f**) the XGB model.

RExPRT misclassifies only four pathogenic TRs as benign: RFC1, FXN, CNBP, and ATXN10. Interestingly, all four of these TRs are in intronic regions within their respective genes. None of them overlap TAD boundaries, RAD21 binding sites, or open regulatory regions and are not characterized as eTRs. They are also not in 3’UTRs or 5’UTRs, nor in promoters except for CNBP. Their GERP scores are extremely close to 0 (range = 0 – 0.039), but the average GERP score for TRs classified as true positives is 0.67. The average GERP score for true negative TRs is −0.10, and there is a statistically significant difference between the two groups (p = 2.80e-09) (Supplementary Fig. 1). The motifs for each of these TRs are unique within the group of pathogenic TRs in the training dataset but are found in the benign group. All these characteristics could explain why these four pathogenic TRs are misclassified as benign.

### RExPRT maintains a low false positive rate on the testing dataset

Next, we ran RExPRT on our testing dataset to assess its performance. The unseen data was comprised of 10 additional pathogenic TRs and 83 rare, benign TR expansions. We found that RExPRT maintained a low false positive rate (1.08%), as seen above (Fig. 4a) while its recall reduced to 60% (Fig 4a, b). Since this dataset has only 10 pathogenic TRs, we expect a volatile recall rate as each additional TR correctly classified improves the rate by 10%. Since our precision (85.71%) is better than our recall (60%) here, the F1 score drops down to 0.71. The ROC curve maintains a steep slope, but its AUC value drops to 0.90 (Fig 4c). The PRC on the other hand suffers a greater reduction and its AUC value falls to 0.74 (Fig. 4d).

**Figure 4.**
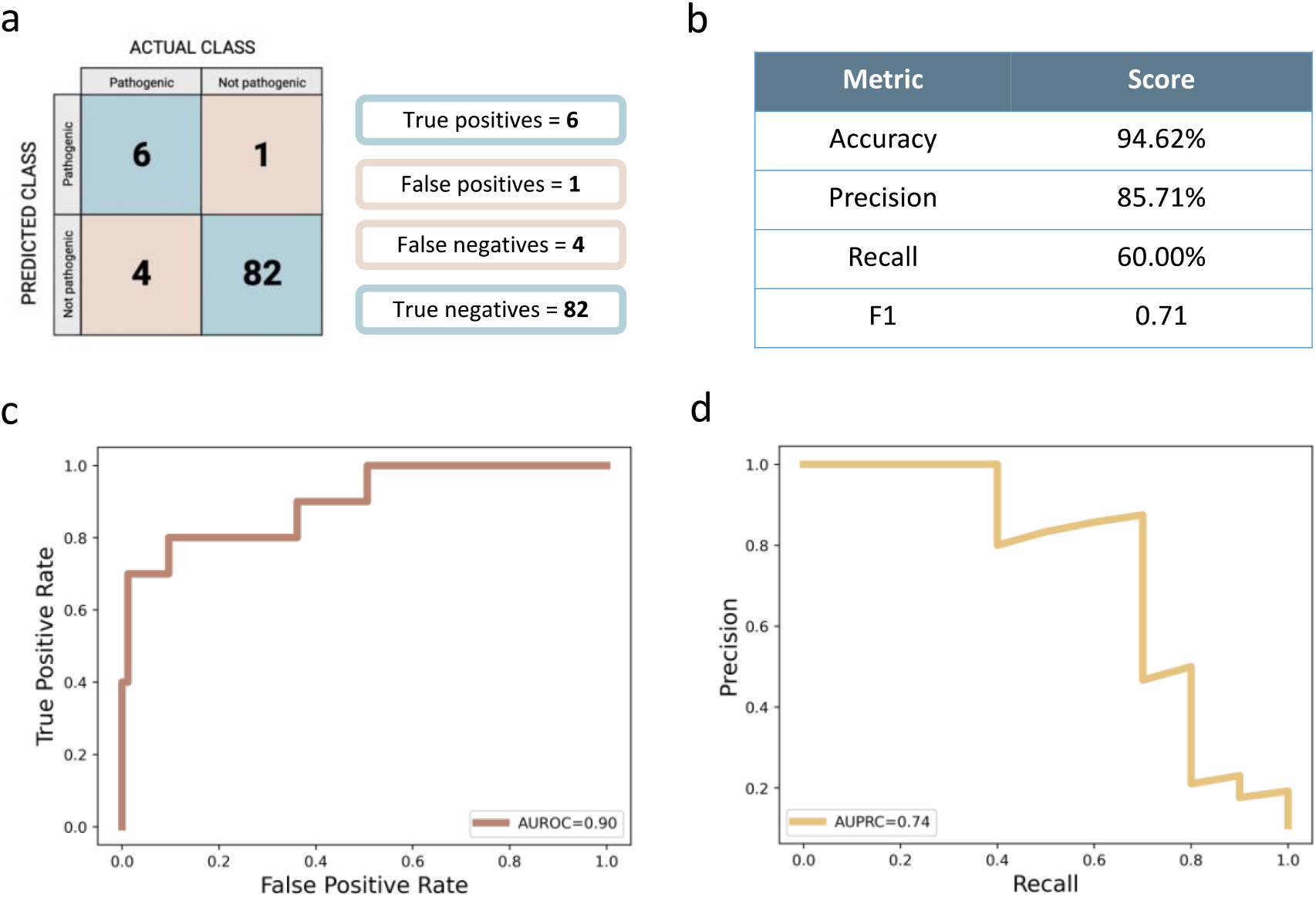
Performance metrics for RExPRT on the testing dataset. (**a**) The confusion matrix results from running RExPRT on the testing dataset of 93 TRs. (**b**) Calculations of the accuracy, precision, recall, and F1 scores. (**c**) ROC curve and its AUC value. (**d**) PRC curve and its AUC value.

There are five pathogenic TRs in the testing dataset whose motifs are a variation of CGG. Since there are 12 such pathogenic TRs in the training dataset, there are ample examples for RExPRT to learn their correct classification. Indeed, RExPRT classified all five of these TRs with CGG motifs as pathogenic. The remaining five pathogenic TRs are all intronic TTTCA expansions in different genes that cause various forms of familial adult myoclonic epilepsy (FAME). There are only two such examples in the training dataset, SAMD12 and DAB1, both of which are correctly classified by RExPRT during LOOCV. In the case of the five FAME loci in the testing dataset, YEATS2 is the only one that is correctly classified as pathogenic, but the XGB score for STARD7 is extremely close to the threshold for pathogenicity with a probability of 0.49. Importantly, these loci are intronic and follow similar patterns in their characteristics as the pathogenic TRs in the training dataset that failed to be classified correctly.

### RExPRT identifies ~30,000 TRs in the reference genome that may be pathogenic if expanded

We ran RExPRT on ~800,000 TRs with motif lengths between 3 and 8bp listed in the hg19 reference genome. This resulted in 29,613 TRs classified as pathogenic if expanded, 67.45% of which are exonic, and 32.53% are intronic (Fig. 5a). Of the benign category, only 0.58% are exonic, and 53.89% are intronic, with the remaining being intergenic (Fig. 5b). Fisher’s exact tests demonstrate a clear enrichment for pathogenic TRs in exons (OR = 53.13; 95% CI = [51.78, 54.36]) (Fig. 5c). Such an observation is not surprising, since expansions in coding regions could alter the structure of the protein. More than half of the TRs predicted as pathogenic are in promoters (24.95%), 3’UTRs (27.56%), or 5’UTRs (13.02%), while only ~3% of the benign TRs fall within these regions (Fig 5d). Fisher’s exact tests confirm an enrichment for pathogenic TRs in promoters (OR = 7.33; 95% CI = [7.13, 7.54]), 3’UTRs (OR = 49.11; 95% CI = [47.65, 50.62]), and 5’UTRs (OR = 45.08; 95% CI = [43.56, 46.74]) (Fig. 5e).

**Figure 5.**
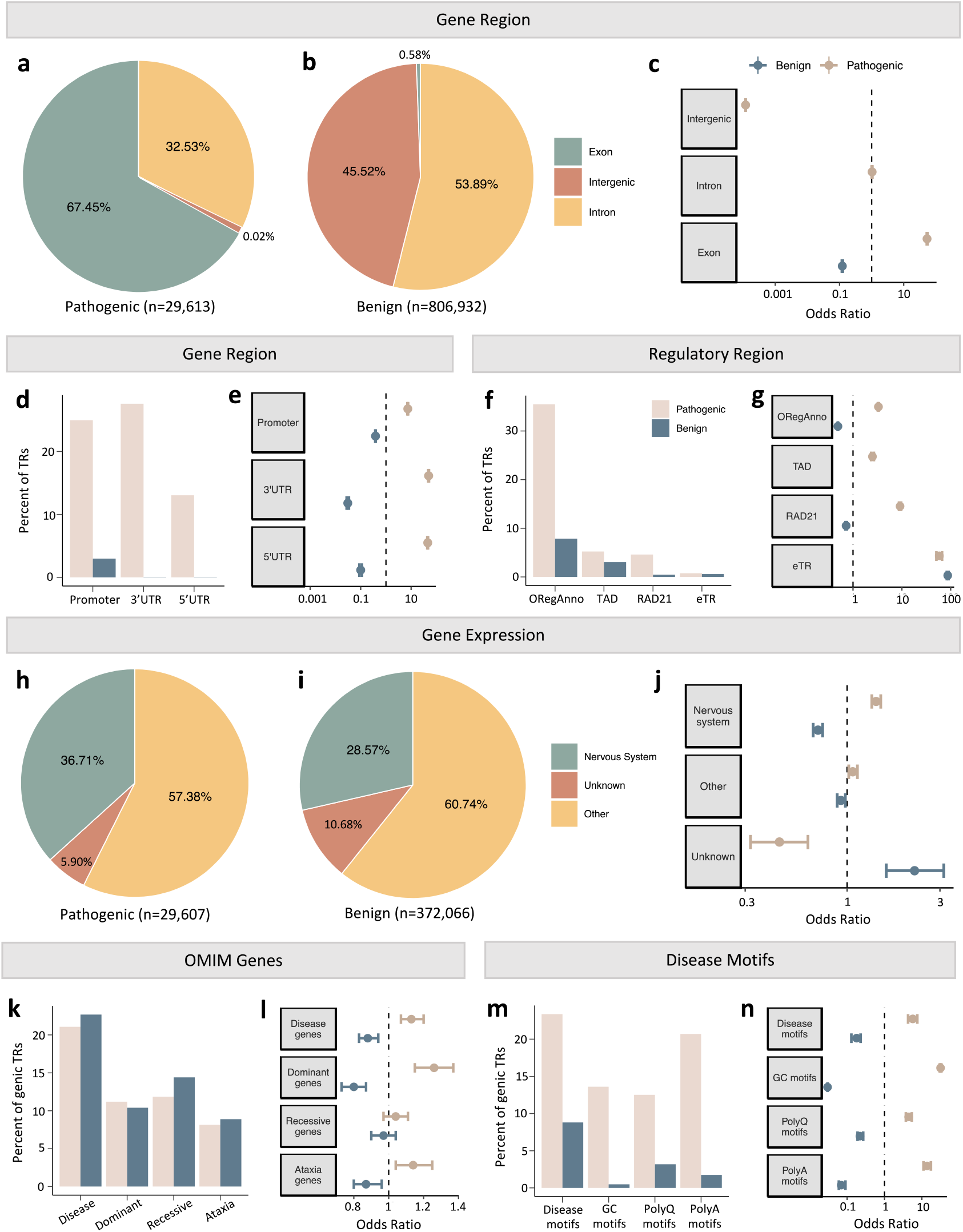
Characterization of ~800,000 reference TRs analyzed by RExPRT. (**a**) TRs in the reference genome that are predicted pathogenic, and their distribution among exonic, intronic, and intergenic categories. (**b**) TRs in the reference genome that are predicted benign, and their distribution among exonic, intronic, and intergenic categories. (**c**) Odd’s ratios for pathogenic and benign TRs in their intersection with exonic, intronic, and intergenic regions of the genome, notably showing an enrichment of pathogenic TRs in exons. (**d**) Distribution of pathogenic and benign TRs in promoters, 3’UTRs, and 5’UTRs. (**e**) Odd’s ratios for pathogenic and benign TRs demonstrating an enrichment of pathogenic TRs in promoters, 3’UTRs, and 5’UTRs. (**f**) Distribution of pathogenic and benign TRs in open regulatory regions, TAD boundaries, RAD21 transcription factor binding sites, and as eTRs. (**g**) Odd’s ratios for pathogenic and benign TRs demonstrating an enrichment of pathogenic TRs in open regulatory regions, TAD boundaries, RAD21 transcription factor binding sites, and eTRs. (**h**) TRs in genic regions that are predicted to be pathogenic, and their tissue expression. (**i**) TRs in genic regions that are predicted to be benign, and their tissue expression. (**j**) Odd’s ratios for pathogenic and benign TRs in their tissue distributions demonstrating an enrichment for pathogenic TRs in nervous system tissues. (**k**) Distribution of pathogenic and benign TRs in OMIM disease genes, dominant disease genes, recessive disease genes, and ataxia genes. (**l**) Odd’s ratios for pathogenic and benign TRs demonstrating an enrichment of pathogenic TRs in OMIM disease genes, dominant genes, and ataxia genes. (**m**) Distribution of pathogenic and benign TRs with known repeat expansion disorder disease motifs, pure GC motifs, polyglutamine (polyQ) motifs, and polyalanine (polyA) motifs. (**n**) Odd’s ratios for pathogenic and benign TRs demonstrating an enrichment of pathogenic TRs with known repeat expansion disease motifs, pure GC motifs, polyQ motifs, and polyA motifs.

We find a greater number of pathogenic TRs overlapping open regulatory regions (35.47%), TAD boundaries (5.22%), RAD21 transcription factor binding sites (4.59%), and being classified as eTRs (0.76%), compared with benign TRs (Fig. 5f). Fisher’s exact tests demonstrate enrichment of pathogenic TRs in TAD boundaries (OR = 2.50; 95% CI = [2.34, 2.66]) and RAD21 binding sites (OR = 9.16; 95% CI [8.71, 9.63]), and in open regulatory regions (OR = 3.30; 95% CI = [3.22, 3.38]). Expression TRs are found in both groups, since these are a subset of the larger group of reference repeats (Fig. 5g).

Next, we removed all intergenic TRs from our pathogenic and benign groups and investigated gene expression. We found that 36.71% of pathogenic TRs are expressed in nervous system tissues, with 57.38% being expressed in other tissues (Fig. 5h). In the benign category, 28.57% are expressed in nervous system tissues, and 60.74% in other tissues (Fig. 5i). Fisher’s exact tests demonstrated an enrichment of pathogenic TRs in nervous system tissues (OR = 1.41; 95% CI = [1.34, 1.49]) (Fig. 5j).

Pathogenic TRs occur at similar rates compared to benign TRs in OMIM disease genes (21.08%), dominant disease genes (11.20%), recessive disease genes (11.84%), and ataxia genes (8.14%) (Fig. 5k). However, Fisher’s exact tests demonstrate a slight enrichment of pathogenic TRs in OMIM disease genes (OR = 1.13; 95% CI = [1.07, 1.20]), as well as in dominant genes specifically (OR = 1.26; 95% CI = [1.15, 1.37]) and ataxia genes (OR = 1.14; 95% CI = [1.04, 1.25]), but not in recessive genes (OR = 1.04; 95% CI = [0.97, 1.11]) (Fig. 5l).

Upon exploring the motifs for TRs in each group, we found that pathogenic TRs have a greater percentage of known repeat expansion disease motifs (23.37%), GC motifs (13.61%), polyQ motifs (12.52%), and polyA motifs (20.71%) compared to benign TRs (Fig. 5m). Fisher’s exact tests demonstrated that pathogenic TRs are enriched in disease motifs (OR = 5.45; 95% CI = [4.12, 7.522), pure GC motifs (OR = 32.56; 95% CI = [31.08, 34.10]), polyQ motifs (OR = 4.35; 95% CI = [3.63, 5.24]), and polyA motifs (OR = 14.83; 95% CI = [11.73, 18.99]) (Fig. 5n).

### RExPRT identifies several interesting candidate genes in the Undiagnosed Diseases Network (UDN) cohort data

We processed 2,982 genomes from the UDN through the outlier pipeline described in Methods that detects rare, repeat expansions in each case. This resulted in 449 candidate TRs, or rare, genic TRs that originate from a reference repeat locus. Of these, 23 genes had TRs that were predicted as pathogenic by RExPRT. Ten of these are already known pathogenic sites, which leaves a total of 13 novel strong candidates for further investigation. More specifically, only three of these are observed in multiple affected patients in a heterozygous state; FAM193B, FRA10AC1, and CLEC2B.

Interestingly, the FAM193B candidate was found in two affected siblings, both with oculopharyngodistal myopathy (OPDM). Notably, this disorder has already been linked to expansions in four other genes: LRP12,^30^ GIPC1,^31^ NOTCH2NLC,^32^ and RILPL1.^33^ All these expansions are composed of a CGG motif and occur in the 5’UTR of their respective gene. The same pattern is observed with the TR we found in FAM193B, making it a strong novel candidate variant for the phenotype. The heterozygous expansion was confirmed in the affected siblings using Oxford Nanopore long-read sequencing, but we are lacking confirmation in additional families. Further investigation into the pathogenicity and mechanism of disease of the expanded TR is underway.

The FRA10AC1 locus has been previously described as likely benign for a phenotype of intellectual disability. The authors concluded that the expansion may be pathogenic only in a homozygous state since methylation of the CGG repeat was observed in a single affected patient and carrier.^34^ However, it may be premature to draw this conclusion since specific repeat sizes were not measured, nor were methylation levels assessed in a cohort of individuals.

The CLEC2B candidate TR was observed in two patients with unsteady gait. The first patient has been partially diagnosed with hearing loss, but this does not explain their symptoms of hypotonia, seizures, and unsteady gait. The second patient has muscle weakness and atrophy, as well as unsteady gait.

## Discussion

We built RExPRT to address the lack of available tools for assessing the pathogenicity of repeats. Our previous data demonstrated that rare repeat expansions are often found in healthy controls,^8^ which indicates that simply selecting TR expansions by allele frequency is not sufficient to determine pathogenicity. RExPRT incorporates information on the genetic architecture of a TR locus, such as its proximity to regulatory regions, TAD boundaries, and evolutionary constraints. It further includes information on gene expression and the DNA motif a TR is composed of. These features enable RExPRT to predict TR pathogenicity with an accuracy of 96.87%. RExPRT excels at having a robust precision due to its low false positive rate. However, its limitations lie in its recall which averages at 75%. In the field of Mendelian discovery genetics, the optimization of precision is preferable to avoid a long list of candidates with many false positive TR expansions that require costly, time-consuming, and labor intensive experimental validation.

Nonetheless, future versions of RExPRT should focus on improving the recall of TRs that are intronic, require motif changes, and are recessive. A major limitation comes from the general paucity of pathogenic TRs to train on, particularly TRs that are recessive, intronic, and require motif changes. By providing the machine learning algorithms with a more comprehensive set of examples to learn from, its performance will likely increase. Additionally, the incorporation of features that are more informative for these problematic TRs could be considered. Finally, since different loci will have different size thresholds for pathogenicity, integration of more accurate genotyping data from long-read sequencing technology may enable future versions of RExPRT to suggest minimum thresholds.

Applying RExPRT to ~800,000 reference TR loci, we found that ~30,000 were classified as pathogenic. It is important to note here that realistically, many of these TRs will never be observed as expanded because they are in fact stable for molecular reasons or affect essential genes or genomic regions that are embryonic lethal. Our results demonstrate that pathogenic TRs, according to RExPRT, are enriched in exons, promoters, 3’UTRs, 5’UTRs, TAD boundaries, RAD21 binding sites, eTRs, and genes expressed in the nervous system. Pathogenic TRs are also enriched in disease genes, particularly dominantly inherited genes, and ataxia genes. Considering many repeat expansion diseases present with ataxia as a primary phenotype, this is a particularly interesting observation. Not surprisingly, our predicted pathogenic TRs are enriched in known disease motifs, including polyQ and polyA. If these TRs are found to be expanded in patients with phenotypes similar to known diseases with the same motifs, they would be extremely strong candidates. FAM193B is one such example; we found the CGG expansion in a family with OPMD, a disease already associated with four other CGG expansions in different genes.^30–33^ Pure GC rich motifs seem to be particularly enriched in the pathogenic group, even in the training and testing datasets. These regions may be associated with mechanisms that make them pathogenic upon expansion.

In conclusion, RExPRT has established the possibility of using machine learning to classify repeat expansion variation. Our work lays the groundwork for similar tools that can be built in the future, mitigating some of its limitations.

## Data availability

The data used in this paper were sourced from our previous publication in *Scientific Data*,^8^ as well as those mentioned in Supplementary Table 1. Additionally, we used dataset EGAD00001003562 available through the European Genome-Phenome Archive.

## Code availability

The scripts to run RExPRT for any TR of interest, as well as the RExPRT scores for the reference TRs are available on GitHub at https://github.com/ZuchnerLab/RExPRT. We are also working towards implementation of RExPRT into GENESIS, a user-friendly point and click analysis tool.

