## Supplementary Materials for "RExPRT: a machine learning tool to predict pathogenicity of tandem repeat loci"

| Feature | Data source | Significance |
|---|---|---|
| **3'UTR** | UCSC annotation | Significant |
| **5'UTR** | UCSC annotation | Significant |
| **Alu elements** | HOMER annotation files | Nonsignificant |
| **Cerebellum eQTL** | UCSC table browser | Nonsignificant |
| **Conserved distal enhancers** | UCSC table browser | Nonsignificant |
| **Constrained non-conserved regions (CNCRs)** | Chen et al[1] | Nonsignificant |
| **CpG islands** | UCSC table browser | Nonsignificant |
| **CTCF (neural cells)** | UCSC table browser | Nonsignificant |
| **EP300 (neural cells)** | UCSC table browser | Nonsignificant |
| **eTR** | Fotsing et al[2] | Significant |
| **Exon** | UCSC annotation | Significant |
| **FAIRE (frontal cortex)** | UCSC table browser | Nonsignificant |
| **H3K27ac (ES cells)** | UCSC table browser | Nonsignificant |
| **H3K4me3 (neurons)** | UCSC table browser | Nonsignificant |
| **Intron** | UCSC annotation | Nonsignificant |
| **LINE** | HOMER annotation files | Nonsignificant |
| **MXI1 (neural cells)** | UCSC table browser | Nonsignificant |
| **ncRNA** | HOMER annotation files | Nonsignificant |
| **Non constrained non-conserved regions (NCNCRs)** | Chen et al[1] | Nonsignificant |
| **ORegAnno** | UCSC table browser | Significant |
| **Promoter** | UCSC annotation | Significant |
| **Pseudogene** | HOMER annotation files | Nonsignificant |
| **RAD21 (neural cells)** | UCSC table browser | Significant |
| **SINE** | HOMER annotation files | Nonsignificant |
| **SMC3 (neural cells)** | UCSC table browser | Significant |
| **TAD boundaries (cortical cells)** | Sun et al[3] | Nonsignificant |
| **TAD boundaries (DLPFC cells)** | PsychENCODE | Significant |
| **TAD boundaries (ES cells)** | Sun et al[3] | Nonsignificant |
| **TAD boundaries with high CpG (DLPFC cells)** | PsychENCODE/UCSC table browser | Nonsignificant |
| **TAD boundaries with low CpG (DLPFC cells)** | PsychENCODE/UCSC table browser | Nonsignificant |

**Supplementary Table 1 Features tested for significant association with pathogenic TRs**

The source where the feature dataset was downloaded from is provided, as well as whether the feature demonstrated significance after conducting fisher's exact tests.

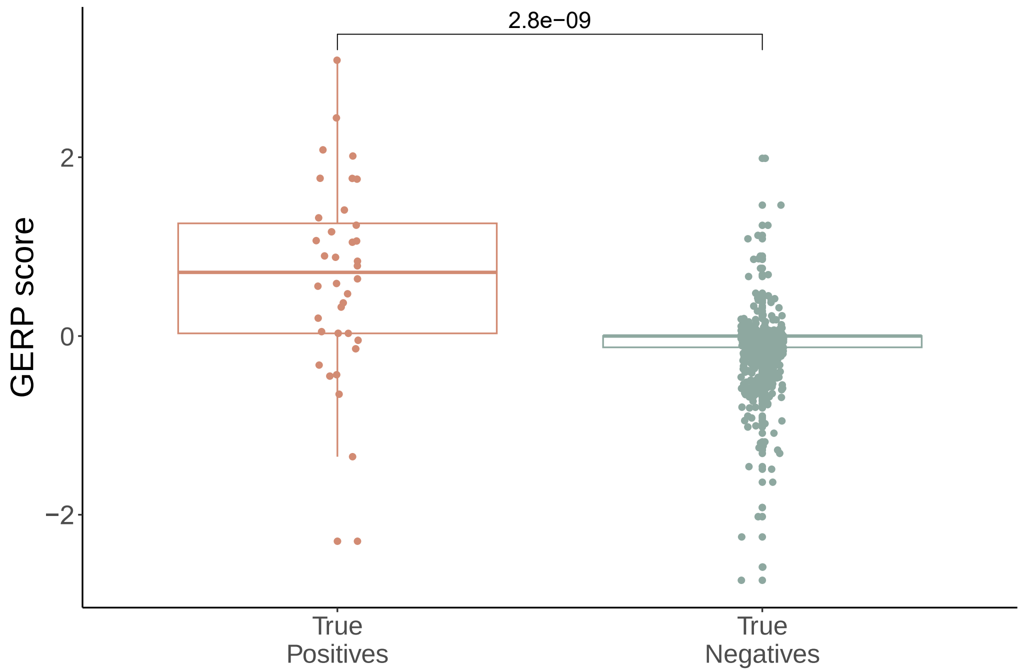


**Supplementary Figure 1: GERP scores for TRs in the training dataset**

GERP scores are plotted for TRs classified as “true positives” or “true negatives” by the RExPRT ensemble method. Wilcoxon signed-rank test p-value demonstrates a statistically significant difference between the two groups.
